# Single-cell RNA-seq analysis maps the development of human fetal retina

**DOI:** 10.1101/423830

**Authors:** Yufeng Lu, Wenyang Yi, Qian Wu, Suijuan Zhong, Zhentao Zuo, Fangqi Zhao, Mei Zhang, Nicole Tsai, Yan Zhuo, Sheng He, Jun Zhang, Xin Duan, Xiaoqun Wang, Tian Xue

**Affiliations:** State Key Laboratory of Brain and Cognitive Science, Institute of Biophysics, Chinese Academy of Sciences, Beijing 100101, China; Hefei National Laboratory for Physical Sciences at the Microscale, Neurodegenerative Disorder Research Center, CAS Key Laboratory of Brain Function and Disease, School of Life Sciences, University of Science and Technology of China, Hefei 230026, China; University of Chinese Academy of Sciences, Beijing 100049, China; Obstetrics and Gynecology Medical Center of Severe Cardiovascular of Beijing Anzhen Hospital, Capital Medical University, Beijing 100029, China; Departments of Ophthalmology and Physiology, Weill Institute for Neurosciences, University of California San Francisco, San Francisco, CA 94143, USA; CAS Center for Excellence in Brain Science and Intelligence Technology, Chinese Academy of Sciences, Shanghai 200031, China; Beijing Institute for Brain Disorders, Beijing 100069, China; Institute for Stem Cell and Regeneration, Chinese Academy of Sciences, Beijing 100101, China

**Keywords:** human retinal development, single-cell transcriptome, retinal progenitor cells, Müller glia, retinal cell lineage

## Abstract

Vision starts with image formation at the retina, which contains diverse neuronal cell types that extract, process, and relay visual information to higher order processing centers in the brain. Though there has been steady progress in defining retinal cell types, very little is known about retinal development in humans, which starts well before birth. In this study, we performed transcriptomic profiling of developing human fetal retina from gestational weeks 12 to 27 using single-cell RNA-seq (scRNA-seq) and used pseudotime analysis to reconstruct the developmental trajectories of retinogenesis. Our analysis reveals transcriptional programs driving differentiation down four different cell types and suggests that Müller glia (MG) can serve as embryonic progenitors in early retinal development. In addition, we also show that transcriptional differences separate retinal progenitor cells (RPCs) into distinct subtypes and use this information to reconstruct RPC developmental trajectories and cell fate. Our results support a hierarchical program of differentiation governing cell-type diversity in the developing human retina. In summary, our work details comprehensive molecular classification of retinal cells, reconstructs their relationships, and paves the way for future mechanistic studies on the impact of gene regulation upon human retinogenesis.

## Introduction

Vision begins at the retina, a sensory laminated structure that converts incoming light into electrical signals and relays these processed signals to the brain^1^. The cell types found within the retina are conserved in vertebrates and generally include six types of neurons (rod and cone photoreceptors, bipolar, amacrine, horizontal and ganglion cells) and Müller glia, all of which are generated from multipotent retinal progenitor cells (RPCs) ^2-5^. Retinogenesis is controlled by the complex interaction of the intrinsic and extrinsic factors upon driving RPCs differentiated into specific cell types ^3,4,6-20^. The generation order of retinal cells has been carefully examined in embryonic rodent retinae using multiple methods, indicating that retinal ganglion cells (RGCs) generate first, following by the generation of horizontal cells (HCs), amacrine cells (ACs), cone photoreceptors, rod photoreceptors, bipolar cells (BCs), and Müller glia (MG) produced last ^4,5,21-24^.

Tremendous progress has been made in uncovering the complex molecular mechanisms that control retinal cell diversification and functions in rodents ^11,25-38^. Several transcriptional factors such as *Nrl*, *Crx*, have been characterized as key regulators for the cell fate determination and classification in retinogensis ^39-43^. However, the studies of cellular and molecular features of retinal cell types are fragmented based on bulk cell analysis. The emerging of single-cell RNA sequencing (scRNA-seq) technologies provides a powerful tool to globally characterize the comprehensive cell types and their molecular characteristics in retinal development ^44-46^. scRNA-seq has been performed to uncover comprehensive subtypes of retinal cells in mouse retina, such as BCs, RGs, and provide insights on the transcriptional network controlling the cell classification and functions ^45,47,48^. However, the scRNA-seq study on human retinal development is blank. Various retinal diseases with vision impairment are associated with abnormal fetal retinal development ^49-51^. The clinical diagnosis and therapy were hampered by limited understanding of the molecular and cellular bases of human retinal development. The precise order of cell genesis, lineage relationships and molecular characteristics has remained poorly understood in the human fetal retina due to the limit access to human retinal samples.

To dissect the cellular and molecular changes in developing human retina, here we utilized scRNA-seq to analyze the transcriptomic features in various developmental stages of human fetal retina. This is the first time that the developing human retina was analyzed by scRNA-seq, and pseudotime analysis reconstructs their developmental trajectories. We identified 34 subtypes of cells within 9 main classes and depicted the developmental trajectory of these cells. We found Müller glia (MG), possible progenies of retinal progenitor cells (RPCs) and appearing as early as GW12 by cell-lineage analysis, are active in proliferation. The lineage analysis based on single-cell transcriptom also suggests that the MG highly likely have the ability to differentiate into neuronal retinal cell types in nature development of human retina. Detailed analysis of RPCs revealed their developmental features. The differentially expressed genes of different subclusters of RPCs were correlated with mature retinal neuron types, suggesting that RPC cell fate is likely pre-determined to generate specific progeny early in retinal development. Our study uncovered the comprehensive molecular classification of retinal cells, reconstructs their relationships, and provide a data sources for future mechanistic studies for human retinogenesis.

## Results

### Cell types of the developing human retina

One or two entire human fetal retinae of gestational weeks 12-27 (GW 12, 14, 16, 18, 22, 24 and 27; 2 biological replicates of GW 24) were analyzed. Droplet-based single cell RNA-Sequencing (10x Genomics Chromium) was performed, resulting in 25,296 individual transcriptional profiles, ranging from 2153 to 4847 cells for each sample, with a median of 1511 genes detected per cell (Supplementary Fig. 1a,b). To classify cell types in the developing human retina, we performed t-distributed stochastic neighbor embedding (tSNE) analysis by Seurat and identified 34 clusters, showing in 2D or 3D views (Fig. 1a,b, Supplementary Fig. 1c). Biological replicates of GW 24 were distributed evenly on the tSNE plot (Supplementary Fig. 1d), indicating that batch effects did not strongly affect clusters. Next, we used known cell-type specific markers to classify these 34 clusters into 9 major cell types, including retinal progenitor cells (RPC, SOX2+ RLBP1- or HES1+ RLBP1-) ^25,26,52,53^, Müller glia (MG), ganglion cells (GCs), amacrine cells (ACs), horizontal cells (HCs), photoreceptor cells (PCs), bipolar cells (BCs), astrocytes and microglia (Fig. 1b,c). To further confirm the characteristics of these cell clusters, we analyzed the gene ontology (GO) of the differentially expressed genes (DEGs) among these 9 major classes (Fig. 1d). We found that RPCs highly expressed genes enriched in cell proliferation while other cell types were enriched for genes in differentiation and functional development. We also enlisted new molecular markers by DEG analysis between the clusters (Fig. 1e, red labels, Supplementary Fig. 1e). Thus, our study reveals the diversity of retinal cells in human fetal retina and their transcriptomic features.

**Figure 1.**
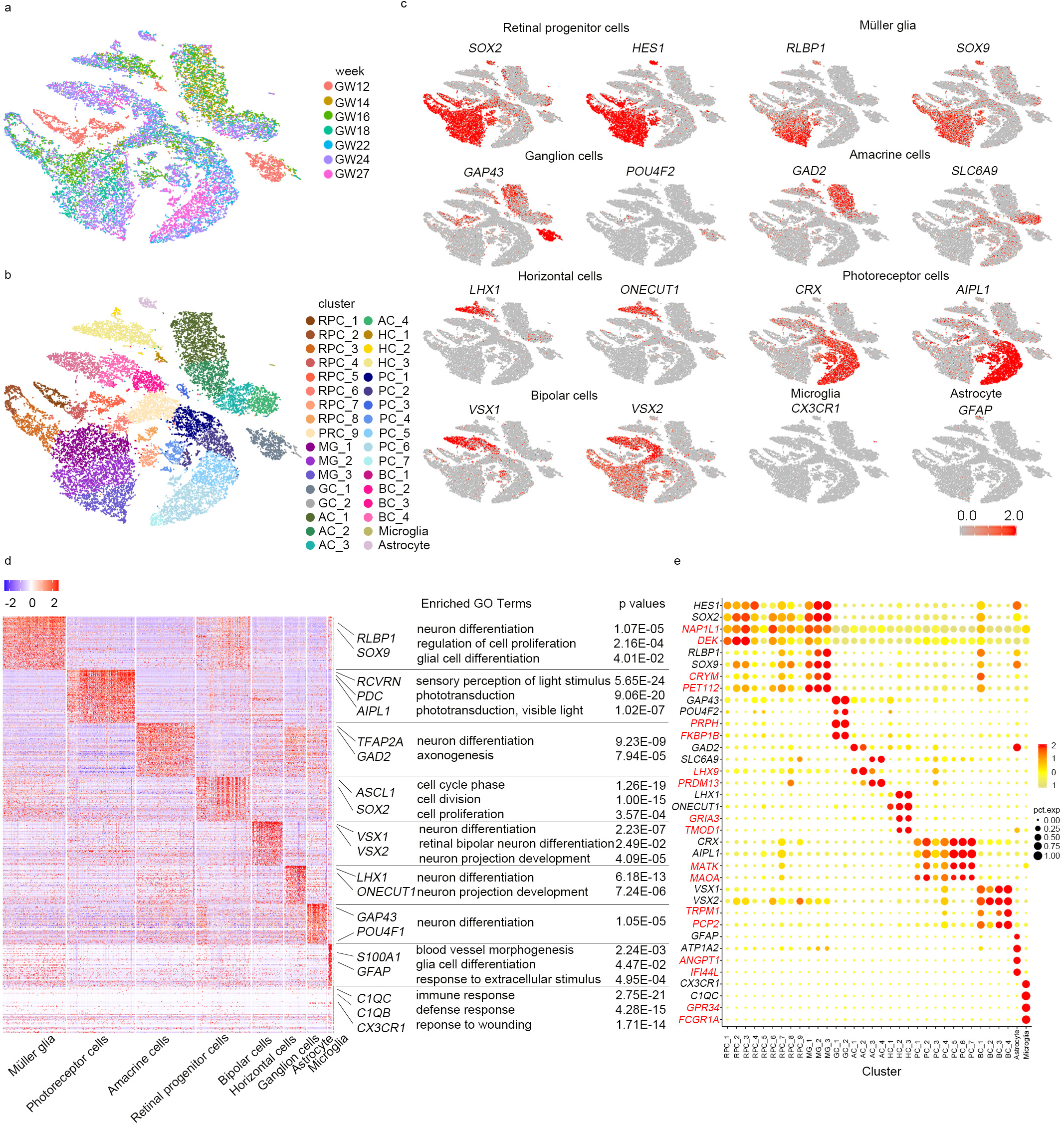
Single-cell Transcriptome Landscape of the Developing Human Retina. a. t-SNE plot of single cells from human fetal retina labelled from GW12 to GW27.
b. Clustering of 25296 single-cell expression profiles into 34 retinal cell populations and 2D visualization of single-cell clusters using t-SNE. Each dot represents a single cell, and clusters are color-coded. (RPC, retinal progenitor cell; MG, Müller glia; GC, ganglion cell; AC, amacrine cell; HC, horizontal cell; PC, photoreceptor cell; BC, bipolar cell.)
c. Expression patterns of known markers for different cell types displayed on t-SNE plots (grey, no expression; red, increased relative expression).
d. Heatmap of blocks of genes enriched within each cell type; specific genes related to each type are highlighted on the right with enriched GO terms. The color key from blue to red indicates low to high gene expression respectively.
e. Dot plot for novel markers of different cell types in the human embryonic retina. Dot plot shows genes enriched within each cluster of cells. Novel markers of each cell type are in red while known markers are in black. The color of each dot (grey, no expression; yellow to red, increased relative expression) shows average scale expression, and its size represents the percentage of cells in the cluster.

### Cell lineages in the developing human retina

Next, we reconstructed the developmental trajectories of single cells in the developing human retina with Monocle ^54-56^. Microglia and astrocytes were excluded from this analysis because microglia are of mesodermal origin and astrocytes are of ectodermal origin ^57,58^. 24979 cells were distributed along a pseudo temporally ordered path that branched into four ends (Fig. 2a, Supplementary Fig. 2a). RPCs and a group of MG were found at the start branch of the developmental trajectory. Cells at End 1 after the first branch point were MG and RPCs, while the rest were progenitors that had differentiated into neuronal cells, including PCs (End 3/4), BCs (End 4) and a mixture of ACs, GCs and HCs (End 2) (Fig. 2a). To further dissect cell lineage, we performed Monocle analysis with the ACs, GCs, HCs and related RPCs. In this analysis, the cell fates of these differentiated cells were distinguished (Fig. 2b). To investigate the transcriptomic differences of cells separated after the start point, moving to End1 or End2/3/4, we then carried out GO analysis on the DEG between three groups of cells, including cells at the start point (RPCs) and the two subgroups divided by the first branching point. The data suggested that the cells on End 2/3/4 undergo visual-related neuronal development, while the cells on End 1 are still active in cell proliferation (Fig. 2c). In addition, we found that the expression of MG markers *RLBP1*, *SLC1A3* and *SOX9* was increased in cells of the End1 branch (Fig. 2d, lines with pink shade) but decreased in cells of neuronal lineage (End2/3/4) (Fig. 2d, lines with blue shade), suggesting that MG are the major cell type of End1. Other than MG, a group of RPCs was also located at End1; hence, the expression of classical progenitor markers, such as *ASCL1*, *HES1*, *SOX2* and *PAX6* decreased in cells of neuronal lineage but sustained high expression in End1 (Fig. 2d). Since RPCs and MG were all located at the start branch of development trajectory with RPCs at the very beginning (Fig. 2a), we further analyzed the lineage of RPCs and MG. Monocle indicated that a group of RPC is the origin of MG in developing human retina. Together, the Monocle analysis suggested that ACs, GCs and HCs are more lineage-correlated but still distinguishable in cell fate, while BCs and PCs are closer in lineage. Interestingly, our data reveal a close relationship between MG and RPCs, indicating the progenitor function of MG in human fetal retina development.

**Figure 2.**
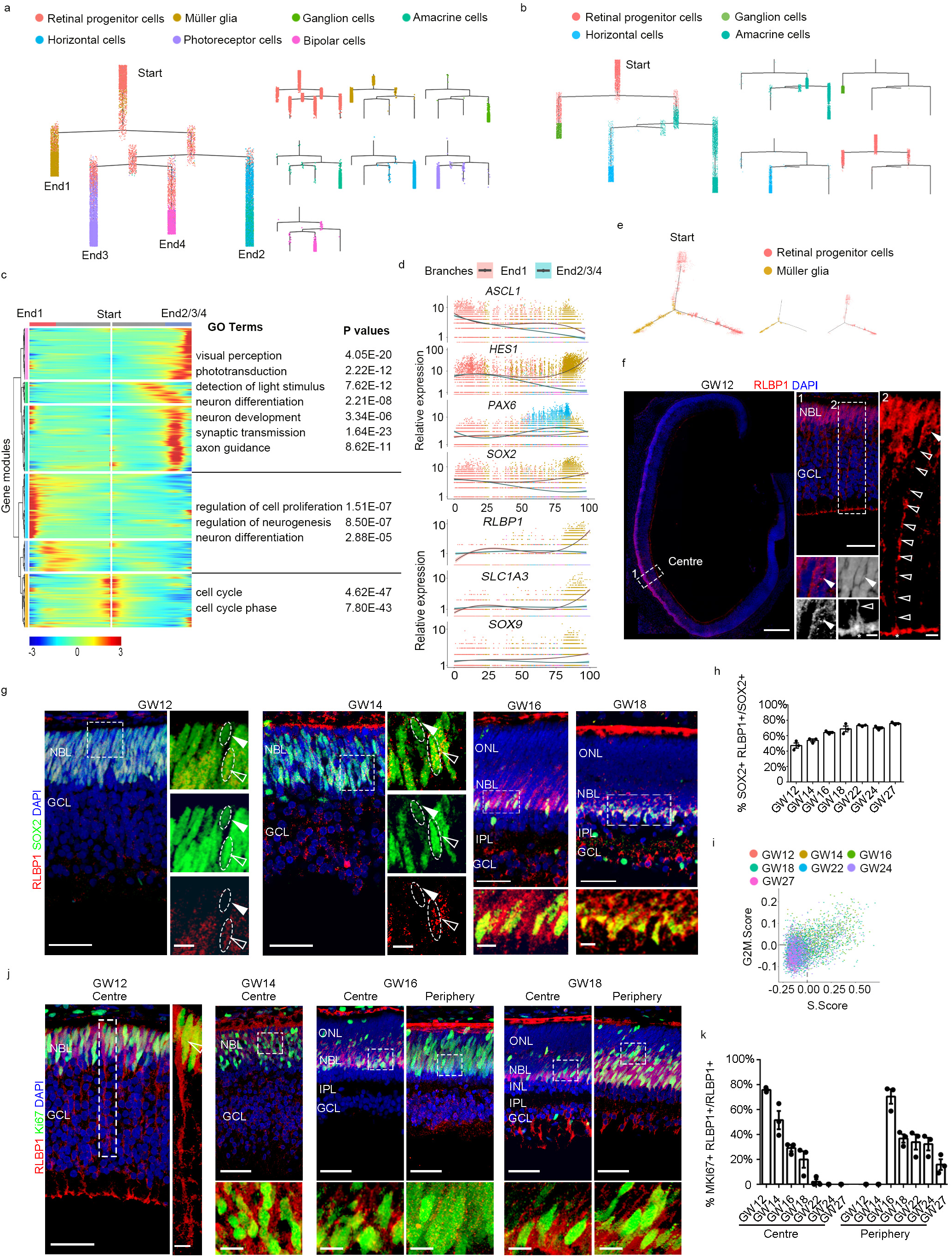
Müller Glia are Embryonic Progenitors in the Human Fetal Retina. a. Cell lineage relationships of all cell types except for microglia and astrocyte in the human embryonic retina. Monocle reveals a branched single-cell trajectory beginning with retinal progenitor cells and terminating at neurons. Each dot represents a single cell; the color of each dot indicates the cell type.
b. Cell lineage relationships of related retina progenitor cells and amacrine cells, horizontal cell, ganglion cells. Monocle recovers a branched single-cell trajectory beginning with retinal progenitor cells and terminating at different cell types. Each dot represents a single cell; the color of each dot indicates the cell type.
c. Gene ontology analysis of modules created by clustering the two main branches from the lineage tree. The analysis reflects the cell fate commitment. In this heatmap, the middle represents the start of pseudo-time. One lineage starts from the start and moves to the End1 while the other lineage moves to the End2/3/4. Rows are genes correlated into different modules.
d. Expression of known markers for retinal progenitor cells and Müller glia. The markers were ordered by Monocle analysis in pseudo-time. The line with pink shadow represents the End1 while blue shadow represents the End2/3/4. The shadow represents the confidence interval around the fitted curve.
e. Cell lineage relationships of retina progenitor cells and müller glia. Monocle recovers a branched single-cell trajectory beginning with retinal progenitor cells and terminating at. Each dot represents a single cell; the color of each dot indicates the cell type.
f. Immunostaining with Müller glia marker RLBP1(red) at GW12 showing the position and morphology of Müller glia in the human embryonic retina. The middle panel is a higher-magnification view of the region boxed in left-most panel and the right-most panel is a higher-magnification view of the regions boxed in middle figure. The four bottom panels in the middle are higher-magnification views of right panel. Arrowheads indicate soma and nuclear; empty arrowheads indicate Müller glia fiber; asterisks indicate Müller glia end foot. Scale bar, 300 μm(left), 50 μm(middle).10um(right),5 μm (bottom 4 figure). Immunostaining of RLBP1 at other gestational weeks are provided in Supplementary Fig.3.
g. Human embryonic retina of GW12, GW14, GW16 and GW18 immunolabeled with SOX2 (green, retinal progenitor cells), RLBP1 (red, Müller glia), showing that Müller glia also express retinal progenitor cell markers. Higher-magnification views of the boxed regions are located to the left or at the bottom of the larger panels. The arrowheads and dotted circle show double positive cells and single positive cells, arrowheads indicate single positive cell, empty arrowheads indicate double positive cells. Scale bar, 50 μm (left), 10 μm (right). Scale bar, 50 um (top), 10 um (bottom). Other gestational weeks of the retina were immunolabeled with SOX2 and RLBP1 as shown in Supplementary Fig.3.
h. Bar charts showing the proportion of RLBP1+SOX2+ cells out of SOX2+ cells in the human embryonic retina at GW12, GW14, GW16, GW18, GW22, GW24 and GW27. Data are presented as mean ± SEM.
i. Cell cycle plot for Müller glia. Each dot represents a single cell, and its color represents the gestational week. The positions of dots illustrate the cell cycle phases of single cells.
j. Immunostaining of Ki67 (green, mitotic cells), RLBP1 (red, Müller glia), showing that some Müller glia can proliferate. Empty arrowheads indicate double positive cells. Scale bar, 50 μm (top); 10 μm (bottom). The centre and periphery of human embryo retina were immunolabeled with RLBP1 and Ki67 at GW22, GW24, GW27 as shown in Supplementary Fig.4.
k. Bar charts showing the proportion of RLBP1+Ki67 + cells in RLBP1+ cells in the centre and periphery of human embryonic retina at different gestational weeks (GW12, GW14, GW16, GW18, GW22, GW24, GW27). Data are mean ± SEM.

### Müller glia are embryonic progenitors

Müller glia are the predominant glial cell type in the retina and serve as support cells for neurons ^59^. In the adult retina, MG bodies are located in the inner nuclear layer (INL) across the entire retina. Among 34 clusters, 3 groups of cells highly expressed *RLBP1*, *SOX9*, *SLC1A3*, *CLU* and *GLUL*, were thus identified as MG (Fig. 1b, c, Supplementary Fig. 2b). Monocle analysis uncovered a group of MG present at the start path of the developmental trajectory, and these MG developed to progenitor (End 1) and neuronal cell fates (End 2/3/4) (Fig. 2a). Transcriptome analysis indicated that MG appear at the entire stage of human retina we obtained, as early as GW12 (Supplementary Fig. 2a), which is in consistent with the evidence that RLBP1+ SOX9+ cells showed up in human developing retina on Day 73 (equals to GW12 based on our development calculation) ^60^. To further reveal the diversity of MG, we used GO analysis of DEGs and found that MG in the early developing retina were enriched in genes for cell proliferation while MG of the late developing stage possessed neuron differentiation genes (Supplementary Fig. 2c). This finding, along with our observation that MG were present at the start path of the Monocle developmental trajectory, led us to investigate whether early born MG might take a role as RPCs. In the wounded adult retina in zebrafish and amphibians, MG can be re-activated to proliferate and differentiate to photoreceptor cells ^61,62^, and human or mouse adult MG can be differentiated into ganglion cell precursors and photoreceptor cells in culture^63-65^. These results suggest the innate properties of MG may be progenitors even in embryonic development. To test this hypothesis, we further analyzed the lineage of the MG sub-clusters using Monocle. The data suggested that the developmental path proceeds from MG-1 to MG-2 to MG-3 (Supplementary Fig. 2d). Consistent with our hypothesis, cells in cluster MG-1/2 were highly enriched with cell proliferation and cell cycle marker genes while MG-3 showing *RCVRN* gene expression, indicating MG may play roles in PC genesis in human retina development (Supplementary Fig. 2c).

To examine the presence and localization of MG in the developing retina, we co-stained RLBP1 and SOX2 in GW8-GW27 retina samples. Interestingly, we found no RLBP1+ cells in GW8 but RLBP1+ cells emerged as early as GW12 in the neuroblastic layer (NBL) of the central retina but not in the periphery where SOX2+ cells are found (Fig. 2f,g, Supplementary Fig. 3a). At GW12 and GW14, the MG arranged into multiple rows in the NBL then gradually moved inwards to finally reside in a single row in the inner nuclear layer (INL). In addition, the MG started to exhibit polarity at GW12 with one side anchored to the inner surface of retina and scaffold structure was observed at GW 24 (Fig. 2g, Supplementary Fig. 3a). The ratio of RLBP1+ SOX2+ / SOX2+ cells in the central retina increased from 47.2% at GW12 to 68.8% at GW18 and was sustained at this level for later times as well (Fig. 2h, Supplementary Fig. 3b).

Since cell proliferation represents a key feature of progenitor cells, we next analyzed the percentages of MG in different stages of the cell cycle (Fig. 2i). Although we found that MG at earlier stages were more mitotically active than ones at later stages, 16.3% and 16.8% of MG in the GW24 and GW27 samples were still in S and G2/M phase respectively (Fig. 2i, Supplementary Fig. 4a). To evaluate this observation, we co-stained developing retina from GW12 to GW27 with RLBP1 and Ki67. Consistent with transcriptome analysis, we observed that a greater number of Ki67+ cells were also RLBP1+ at earlier (GW12) compared to later times (Fig 2h,i, Supplementary Fig. 4b). Interestingly, as MG appeared in the central retina at GW12 and GW14, proliferation of MG in this area also increased in parallel. As the retina develops, the Ki67+ MG decreased in the central region but increased in the periphery (Fig 2j,k, Supplementary Fig. 4b). These observations are consistent with the MG subcluster lineage and GO analysis results (Supplementary Fig. 2b, c), which indicates that some MG may differentiate into neurons. Together, our data supports that MG with proliferative capability serve as RPCs and contribute to embryonic retinogenesis.

### Fate of RPCs is specified at an early time

To further understand the molecular characteristics of RPCs, we analyzed the DEGs of 9 RPC clusters (Fig. 3a). Interestingly, we found that RPCs in Cluster 5 (from GW 12-16) differentially expressed early GC marker *GAP43*, while *AIPL1* and *RECOVERIN* were high in RPCs of Cluster 7, suggesting that these cells may develop to PCs. RPCs in Cluster 8 expressed *TFAP2A* and *TFAP2B*, which are early AC and HC markers. In addition, *VSX2* and *OTX2* were high in RPCs of Cluster 9, indicating that they may develop into BCs and PCs (Fig. 3a). Next, we reconstructed RPC developmental trajectories using Monocle (Fig. 3b, Supplementary Fig. 5a). We observed that one group of RPCs remained as self-renewing progenitors, which we label in the Monocle pseudo-time path as RPCs; while the other groups of RPCs became specified progenitors set to differentiate into functional neurons or MG; we labeled these RPC subtypes as pre-PCs, pre-GCs, pre-ACs, pre-HCs, and pre-MG in the pseudo-time path (Fig. 3b). To further investigate the cell fate of RPCs, we align RPCs that with no Ki67 expression with the differentiated cell types across datasets based on variations as identified by canonical correlation analysis ^66^ (Fig. 3c, Supplementary Fig. 5b). In consistent monocle analysis, we observed RPCs in Cluser 4, Cluster 5, Cluster 7 develop to HCs, GCs and PCs, respectively. RPCs in Cluster 8 show the potential to ACs and HCs differentiation while RPCs in Cluster 9 present cell fate to BCs and PCs. Additionally, we found RPCs in Cluster 3 and Cluster 6 may be the ancestor of MG cells (Fig. 3c). From the trajectories, the pre-MG and a small group of pre-PCs, as well as major pre-GCs, pre-ACs and pre-HCs branch off early in the path, indicating that they specified their identity early on. The group of pre-PCs and pre-MG were closer in lineage, while the pre-GCs, pre-ACs and pre-HCs were clustered in another branch. Major pre-PCs and pre-BCs were generated at a relatively late time (Fig. 3b, Supplementary Fig. 5c). The RPCs were mapped to their developmental trajectory plot by their gene expression profiles, indicating that the determination of RPC cell fate is possibly regulated by the intrinsic factors of RPC, such as expressing various transcriptional factors.

**Figure 3.**
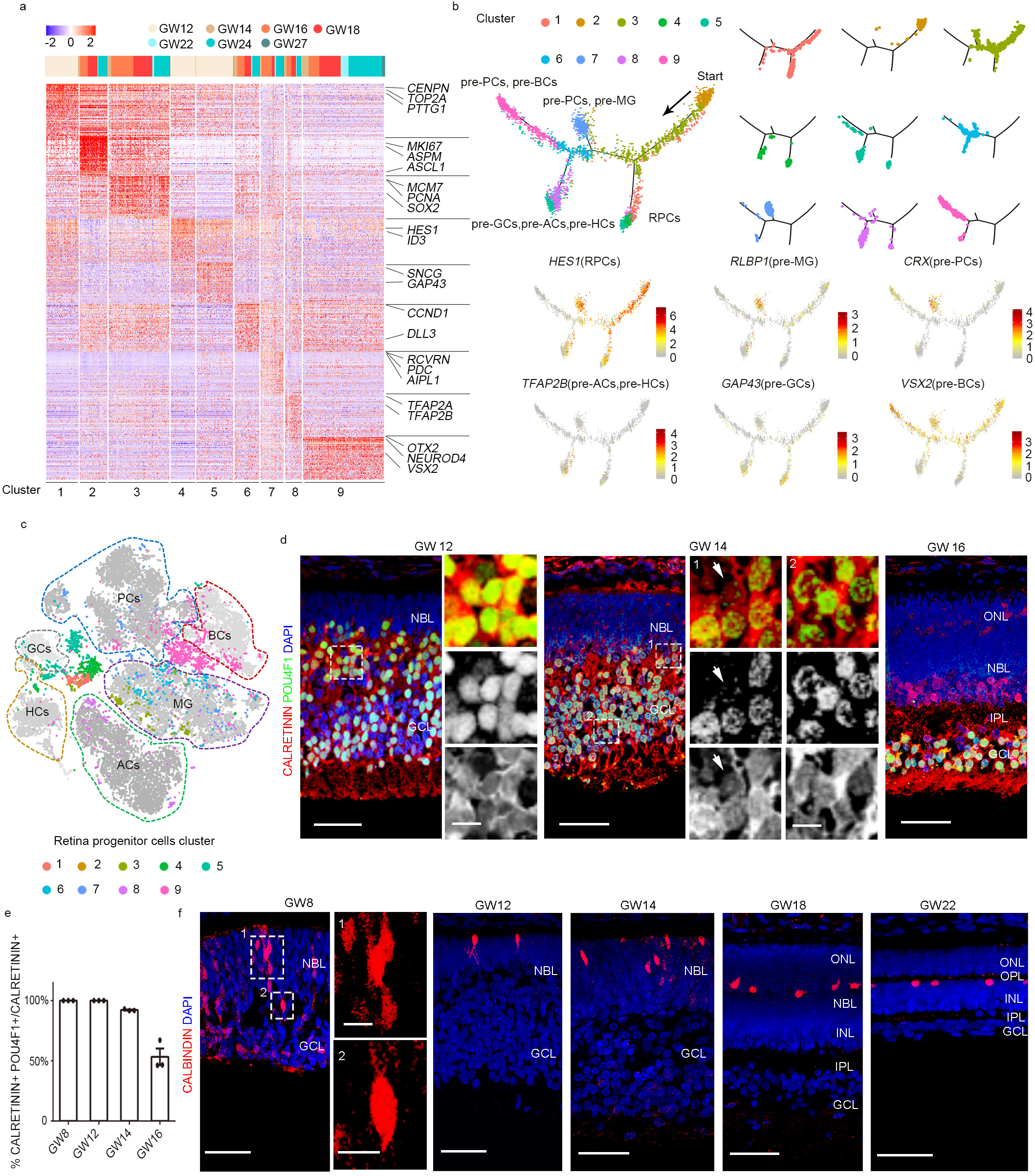
Molecular Features of Neural Progenitor Cells of the Developing Human Retina. a. Heat map showing the gene expression patterns of subclusters of retinal progenitor cells. Known markers for each cell type are shown on the right of each heatmap panel. The graph on the top shows the distribution of each subclass by gestational week. The color key from blue to red indicates low to high expression levels, respectively.
b. Cell lineage relationships of retinal progenitor cells in the human embryonic retina. Monocle recovered a branched single-cell trajectory beginning with retinal progenitor cells and ending with different subtypes of cells with pre-determined developing fate. Each dot is a single cell; the color of each dot represents the cell type. Expression of cell type markers *HES1*(RPCs), *RLBP1*(pre-MG), *CRX*(pre-PCs), *TFAP2B* (pre-ACs, pre-HCs), *GAP43*(pre-GCs), *VSX2*(pre-BCs) showed that the retinal progenitor cells differentiate to different cell types.
c. Integration of retina progenitor cells and different types of retina cells based on variation. The markers of different cell types were shown in Supplementary Fig.5b.
d. Immunostaining with CALRETININ (red, amacrine cells, horizontal cells, ganglion cells) and special ganglion cell marker POU4F1 (green) in the human retina at GW12, GW14 and GW16. At GW12, all the CALRETININ+ cells express POU4F1, showing there are no amacrine cells. The amacrine cells (CALRETININ+/POU4F1-) began to be detected at GW14. At GW16 there are distinct layers for horizontal cells, amacrine cells and ganglion cells (from top to bottom). The arrows show single positive cells (CALRETININ+/POU4F1-). Scale bar, 50 μm; higher-magnification views Scale bar, 10 μm. Immunostaining of CALRETININ in other gestational weeks are shown in Supplementary Fig.6b.
e. Bar charts showing the proportion of CALRETININ+ POU4F1+ cells out of CALRETININ+ cells in the human embryonic retina at GW8, GW12, GW14, GW16. Data are mean ± SEM.
f. CALBINDIN (red, horizontal cells), showing some cells have already acquired horizontal cell fates at GW8. CALBINDIN+ cells show a scattered distribution at GW14 but begin to form a distinct layer at GW18 and have migrated to their appropriate layers by GW22. Immunostaining of GW27 provided in Supplementary Fig.6c.

To validate our *in silico* results on the early generation of GCs, ACs and HCs, we used cell-specific markers to reveal the existence and localization of these cells. CALRETININ and POU4F1 were used to define GCs (CALRETININ+ POU4F1+) and ACs (CALRETININ+ POU4F1-). At GW8, all CALRETININ+ cells were also POU4F1+ and localized in the ganglion cell layer (GCL). However, a few CALRETININ+ POU4F1-ACs appeared at GW14, on top of GCs in the NBL. The number of ACs dramatically increased at GW16 and aligned in the inner part of the INL at GW18 (Fig. 3d,e, Supplementary Fig. 6a,b). CALBINDIN was used as an HC marker, and we observed CALBINDIN+ cells at GW8, mainly localizing in the NBL (Fig. 3f), suggesting that the existence of HC can be detected as early as GW8. At GW12, branches of CALBINDIN+ cells were observed, and HCs migrated from the NBL down to the INL that forms in the retina (Fig. 3f, Supplementary Fig. 6c). These data are consistent with our scRNA-seq analysis and prior reports on the generation of different retinal cell types in the human and mouse retina ^3-5,60^.

### Neuronal cell subtype differentiation and classification

Next, we looked at the molecular differences between sub-clusters of GCs, HCs, ACs and BCs. Cells in Cluster GC-1, the majority of which are from GW12, highly expressed newborn neural markers such as *TUBB3*, while cells in GC-2 were relatively more mature as they expressed several functional neural genes such as SNAP25 (Supplementary Fig. 7a). The limited clusters suggest that GCs may remain to be further differentiated into different subtypes beyond GW27. Cells in HC-1 expressed high levels of *SOX2*, indicating that they were likely specified HC precursors while those in HC-2/3 were mature, fully differentiated cells with high expression of *LHX1*, *ONECUT1* and *PROX1* (Supplementary Fig. 7b). ACs that use GABA as their neurotransmitter (GAD1+ and GAD2+) belonged to Cluster AC-1 and were born earlier than those that use glycine (SLC6A9+) in AC-3/4 (Supplementary Fig. 7c). Cone bipolar cells were classified into BC-3 and BC-4 according to marker genes *VSX1* (Supplementary Fig. 7d).

We next investigated the maturity of the ACs and BCs in our samples. Since many mature subtypes of ACs and BCs have been identified in mice ^45,47^, we wondered if these mature mouse subtypes are also present in the developing human retina (Supplementary Fig. 7e, f). We referenced known subtype markers ^45^ and found a very limited number of ACs and BCs subtypes: *CCK*+ ACs, a type of AC that makes contacts with rod bipolar cells and ON-cone bipolar cells, in Cluster AC-4; a mixture of *PDGFRA*+ and *GBX2*+ ACs in Cluster AC-2; and MAF+ SLITRK6+ ACs in Cluster AC-1 (Supplementary Fig. 7e). In addition, *NNAT*+ and *SEPINI1*+ BCs were found in BC-2 and BC-4 respectively (Supplementary Fig. 7f). These data indicate that although several subtypes of ACs and BCs are generated early in retinal development, the majority of subtypes of ACs and BCs cannot be identified even at GW27 according to the recent references from adult mouse retina studies. Thus, similar to analysis of GCs, these data also suggest that BCs and ACs will likely further specify their subtype properties in later gestational and pediatric stages.

### Developmental features of photoreceptors

Photoreceptors, a specialized type of retinal sensory neuron that is capable of visual phototransduction, were also found to differentiate early from RPCs (Fig. 3a). Hence, we analyzed the 7 clusters of PCs and observed that the cells of Clusters PC-1/2/3/4, and PC-5/6/7 expressed genes related to cell differentiation and visual functions respectively. These two groups also appeared to follow a developmental timeline (Fig. 4a), so we analyzed the GO of genes expressed by PCs from each gestational week. At GW16, axonogenesis may occur in PCs, while genes of response to light stimulus and phototransduction were enriched at GW24 and GW27 (Fig. 4b, Supplementary Fig. 8a), indicating that PCs may be functional at early gestational weeks. To validate the birth of PCs in developing human retina, we stained the retina with RECOVERIN, S-OPSIN, L/M-OPSIN and RHO antibodies, to reveal immature PCs, S cones, L/M cones and rods (Fig. 4c, Supplementary Fig. 8b-e). We found that RECOVERIN+ PCs were accumulated to the most outer layer of NBL as early as GW 12, some of which already developed long protrusions to the inner part of NBL or even to the GCL (Fig. 4c). S cones and L/M cones arose at GW16 and GW22 respectively, and rods also showed up at GW22 with relatively classical shapes in the ONL (Fig. 4c, Supplementary Fig. 8b-e). Both cones and rods appeared mature structure with clear out segment at GW27 (Fig. 4c). To find new markers for PCs, highly expressed genes in Cluster PC-1-7 were selected in the developing retina, including *USP2*, *LAYN*, *PLA2G4C*, *ANXA4*, *DOCK8*, *GPR160*, *MARCH1*, *RRAD*, *TMEM138* and *EPS8* (Fig. 4d, Supplementary Fig. 9). Altogether, this data uncovers the heterogeneity of PCs of the developing retina and reveals new molecular markers.

**Figure 4.**
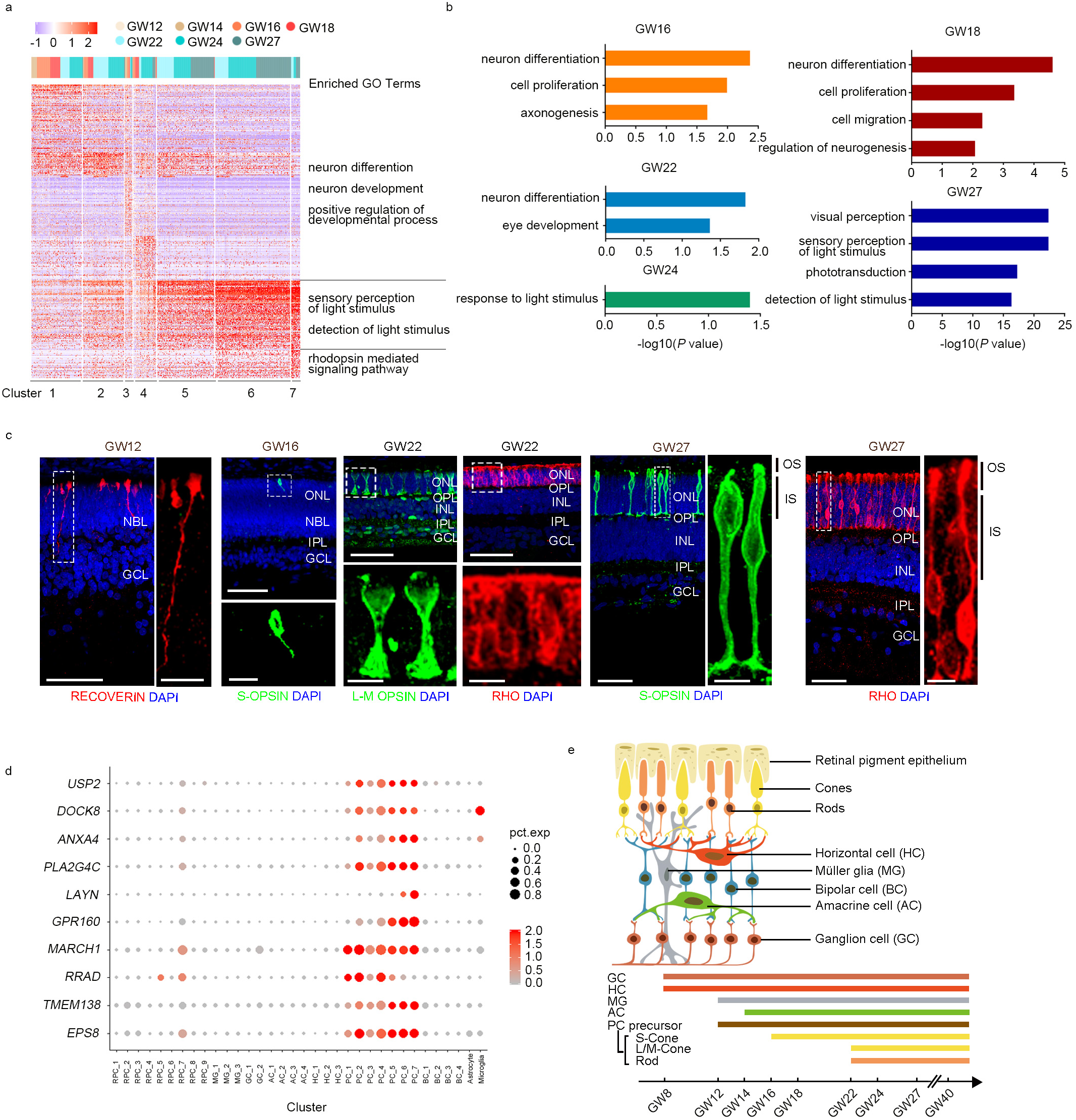
Photoreceptor Cell Development in the Developing Human Retina. a. Heat map of differentially expressed genes of subcluster of photoreceptor cells. The photoreceptor cells go through two developmental stages from GW12 to GW27. The bar chart on the top shows gestational week information. The color key from blue to red indicates low to high gene expression.
b. The enriched gene ontology terms show the cell properties of the photoreceptor cells at different gestational weeks.
c. Immunostaining with RECOVREIN (early photoreceptor), RHO (Rod photoreceptor), S-OPSIN(S-Cone), L/M-OPSIN (L/M-Cone), showing that photoreceptors have begun to develop at GW12, S-Cone appear at GW16, L/M-Cone appear at GW22, and rods appear at GW22. The rod and cone both mature with complete structure at GW27. OS, outer segment; IS, inner segment. Scale bar, 50 μm; higher-magnification views Scale bar, 10 μm. Immunostaining of RECOVERIN, S-OPSIN, L/M-OPSIN, RHO from other gestational weeks is shown in Supplementary Fig.8.
d. Dot plot for novel markers of photoreceptor cells. The size of each dot represents the percentage of cells in each cluster. Grey to red indicates a gradient from low to high gene expression.
e. Schematic representation of major cell types in the three layers of the retina. Color bar represents the developmental birth order of retinal cell types.

## Discussion

In current study, we conducted scRNA-seq in various developmental stages of human retinae to study the cellular and molecular dynamics along retinogenesis. This is the first time that human retinal development has been investigated in time series in single cell resolution. In our study, the analysis of the single cell molecular features uncovered retinal cell types and their birth order, reconstructed the developmental trajectories, and established the cell lineage relationship in human retina. The data also demonstrated that RPCs specified into progeny subclasses which more commit to seven cell types of the retina, followed by the genesis of specific retinal cell types. Interestingly, we detected a subtype of early-born proliferating Müller glia which may function as progenitors in the human fetal retina. Overall, this report provides a data resource for understanding human retinal development in the early and mid-gestational stages (Fig. 4e).

The heterogeneity of RPCs was observed in our study by using scRNA-seq analysis. Multiple subclasses of RPCs were identified in our study based on their distinct gene expression, consistent with their respective differentiated cell types. Previous studies have already demonstrated that there are subclasses of RPCs existed in the mouse retina—the early RPCs and the late RPC ^4,67-69^. The early and the late RPCs in the mouse retina are distinct in gene expression and in their ability to differentiate into glial cells and subtypes of neurons. Here, in the developing human retina, we observed a similar phenomenon that subclasses of RPC are distinct in gene expression and in their ability to differentiate into specific cell types. Therefore, our data provide a source for future molecular study of RPCs diversification and differentiation. Since our data were only collected till GW27 and we still found some RPCs or MG active in cell cycle, we cannot conclude too much beyond this development time on the cell fate determination. Further studies in non-human primate may fill this gap.

We found MG as early as GW12. Consistent with the previous study, MG were detected early than PCs in human retina^60^, whereas in previous studies in mice, MG appeared much later^3-5^. Cell lineage analysis indicated that the MG are from RPCs and we have not observed RLBP1+ cells in GW8 retina samples, indicating that MG may not be the origin of retinogenesis. However, MG were detected in all samples from GW12-27, and the early generation of MG in human retina suggest that MG have additional roles in human retinal development. Indeed, our data analysis suggest MG may play a role as the progenitors. By co-stained RLBP1 with KI67, we found a subpopulation of MG with high proliferation capability which is a unique feature of RPCs. The hypothesis of MG as RPC has been proposed in previous studies^70^. In zebrafish and amphibians, MG have the ability to differentiate into neurons and the ability to self-renew after injury^61,62^. Mammalian MG are able differentiated into neuron after injury with the additional expression of several key factors^63,71-73^. These results suggest the innate properties of adult MG may be progenitors but they are quiescent in mature retina. In GW27 samples, we observed that MG cell body already located in the inner nuclear layer of the retina and they span across all retinal layers, having processes reach and form part of the outer and inner limiting membranes, which exhibits the similar structure as known MG in adult retina. And not many RLBP1^+^ MG have proliferating abilities indicating that these MG may be close to adult MG. Our cell lineage data suggest MG may differentiate to photoreceptor cells in embryonic development, which is in consistent with *in*-*vivo* or *in*-*vitro* MG activation and differentiation.

The developmental lineage relationships of different retinal cell types have been an area of great interest. Our data reconstructed cell lineage relationships in the human retina, showed that MG and RPC are closely related in lineage; ACs, GCs and HCs are more lineage-correlated; some PCs and BCs are more lineage-related. The scRNA-seq data will provide new insights on the gene regulatory networks that account for these unique identities of these cell types. The retina of humans possesses a unique central architecture-fovea centralis, for high acuity vision^74^. Previous study suggests that the central fovea and surrounding macula differentiate much faster than the peripheral retina^75-80^. The dissection and collection of enough fovea cells for 10x scRNA-seq in the developing human retina is challenging in our study, so we did not include any separate fovea data here. While our data collected from the whole human fetal retina, so we did not lose the information in fovea in data sets. Consistent with the previous studies, we indeed detected the development of fovea is faster and earlier than peripheral retina. The positive staining of RLBP1+KI67+ was detected in central retina earlier than peripheral retina, and the staining deceased in central retina following by increased staining signals in peripheral retina. Our further study will focus on the scRNA-seq analysis of fovea region only, and explore its molecular features.

Inherited retinal diseases are a group of retinal disorders caused by an inherited gene mutation and can result in vision loss or blindness^49-51,81^. Currently, the disease diagnosis and therapies were hampered by our understanding the development of human fetal retina. Previous studies have explored using induced pluripotent stem cells (iPSCs) and organoid cultures as stem cell-based approaches for patient-specific treatment for retinal regeneration^82-86^. However, the advancement of stem cell therapy was hampered by our limited knowledge on the molecular mechanisms underlying retinogenesis and disease progression in humans, too. Hence, the transcriptome profiling of individual retinal cells not only pave a way for constructing the development progression of human retina, but also shed lights on understanding the molecular mechanisms underlying retinal degenerative diseases.

## Acknowledgments

This work was supported by the Strategic Priority Research Program of the Chinese Academy of Sciences (XDA16020600), National Key Research and Development Program of China (2017YFA0103303, 2017YFA0102601); this work was made possible in part, by That Many May See pilot fund, NIH-NEI EY002162 - Core Grant for Vision Research and by the Research to Prevent Blindness Unrestricted Grant (X.D.).

## Author contributions

X. W., T. X. and X. D. conceived the project, designed the experiments. Y. L. and S. Z. performed RNA-seq. Y. L., S. Z. and Z. Z. analyzed the data. Y. Z. and S. H. coordinated computational analyses. W. Y. and Q. W. performed immunofluorescence and imaging. F. Z. and J. Z. prepared the samples. Q. W., M. Z. and N. T. wrote the manuscript with input from all other authors. All authors edited and proofread the manuscript.

## Data and materials availability

The single-cell RNA-seq data used in this study have been deposited in the Gene Expression Omnibus (GEO) under the accession number GSE116106.

## Materials and Methods

### Ethics statement

The human embryo collection and research analysis was approved by the Reproductive Study Ethics Committee of Beijing Anzhen Hospital (2014012x). The informed consent forms were designed under ISSCR guidelines for fetal tissue donation and were in strict observance of the legal and institutional ethical regulations for elective pregnancy termination specimens at Beijing Anzhen Hospital. Informed consent for fetal tissue procurement and research was obtained from the patient after her decision to legally terminate her pregnancy but before the abortive procedure. All tissue samples used in this study were not previously involved in any other procedures. All protocols were in compliance with the ‘Interim Measures for the Administration of Human Genetic Resources’ administered by the Chinese Ministry of Health.

### Tissue sample collection and dissociation

Gestational age was measured in weeks from the first day of the woman’s last menstrual cycle to the sample collecting date. Fetal retina tissue samples were collected in ice-cold artificial cerebrospinal fluid (ACSF) containing 125.0 mM NaCl, 26.0 mM NaHCO3, 2.5 mM KCl, 2.0 mM CaCl2, 1.0 mM MgCl2, 1.25 mM NaH2PO4; pH 7.4, bubbled with carbogen (95% O2 and 5% CO2). The samples were gently separated into small pieces and then centrifuged at 200g for 2min. The supernatant was then removed and 500ul digestion buffer (2mg/ml collagenase IV (Gibco), 10 U/μl DNase I (NEB), and 1mg/ml papain (Sigma) in PBS) were added. The tissue was then rotated and incubated at 37°C on a thermo cycler with 300g for 15-20 min. The sample was pipet every 5 min to digest the tissue sample into single cells ^87^. Finally, cessation buffer of equivalent volume (10% fetal bovine serum (Gibco) in PBS) was added to stop digestion.

### Immunohistochemistry

Human retina tissue samples were fixed overnight in 4% paraformaldehyde. The fixed retina was dehydrated in 20% and 30% sucrose in PBS at 4°C and embedded in optimal cutting temperature medium (Thermo Scientific). Thin 20-40μm cryosections were collected on superfrost slides (VWR) using a Leica CM3050S cryostat. For immunohistochemistry, antibodies against the following proteins were used at the indicated dilutions: Goat anti-SOX2 (1:250, Santa Cruz), Rabbit anti-Mki67 (1:200, Chemicon), Rabbit anti-Recoverin (1:1000, Millipore), Mouse anti-POU4F1 (1:500, Santa Cruz), Mouse anti-RLBP1 (1:500, Abcam), Mouse anti-Rod-OPSIN (1:1000, Sigma), Rabbit anti-S-OPSIN (1:500, Millipore), Rabbit anti-L/M-OPSIN (1:500, Millipore), Mouse anti-Calbindin (1:500, Abcam), and Goat anti-Calretinin (1:500, Millipore). Primary antibodies were diluted in blocking buffer containing 10% donkey serum, 0.2% Triton X-100 and 0.2% gelatin in PBS at pH 7.4. Alexa Fluor 488, Alexa Fluor 594 or Alexa Fluor 647 fluorophore-conjugated secondary antibodies (1:500) (Life Technologies) were used as appropriate. Cell nuclei were stained with DAPI (1:10000) (Life Technologies). Images were collected using an Olympus FV1000 confocal microscope.

### Library preparation for high-throughput sequencing

Cells were suspended in 0.04% BSA/PBS and stained with 7-amino-actinomycin D (7-AAD) for 10 min on ice. 7-AAD-negative cells were collected by FACS and re-suspended in 0.04% BSA/PBS at the proper concentration to generate cDNA libraries with Single Cell 3’ Reagent Kits, according to the manufacturer’s instructions. Thousands of cells were partitioned into nanoliter-scale Gel Bead-In-EMulsions (GEMs) by 10x™ GemCode™ Technology, in which the cDNA produced from the same cell share a common 10x Barcode. Upon dissolution of the Single Cell 3’ Gel Bead in a GEM, primers containing an Illumina R1 sequence (read1 sequencing primer), a 16 bp 10x Barcode, a 10 bp randomer and a poly-dT primer sequence were released and mixed with the cell lysate and Master Mix. After incubation of the GEMs, barcoded, full-length cDNA from poly-adenylated mRNA were generated. The GEMs were then broken, and silane magnetic beads were used to remove leftover biochemical reagents and primers. Prior to library construction, enzymatic fragmentation and size selection were used to optimize the cDNA amplicon size. P5, P7, an index sample, and R2 (read 2 primer sequence) were added to each selected cDNA during end repair and adaptor ligation. P5 and P7 primers were used in Illumina bridge amplification of the cDNA (http://10xgenomics.com). Finally, the library was sequenced with 150bp pair-end reads using an Illumina HiSeq4000.

### Data processing of single-cell RNA-seq from Chromium system

Cell ranger 2.0.1(http://10xgenomics.com) was used to perform quality control and read counting of *Ensemble* genes with default parameter (v2.0.1) by mapping to the hg19 human genome. We excluded poor quality cells after the gene-cell data matrix was generated by cell ranger software using the Seurat package (v2.2.0) (https://satijalab.org/seurat/pbmc3k_tutorial.html) in Bioconductor ^45,66,88^. Only cells that expressed more than 850 genes and fewer than 8000 genes were considered, and only genes expressed in at least 3 single cells (0.01% of the data) were included for further analysis. Cells that expressed haemoglobin genes (*HBM*, *HBA1*, *HBA2*, *HBB*, *HBD*, *HBE1*, *HBG1*, *HBG2*, *HBQ1* and *HBZ*) were also excluded. Cells with the mitochondrial gene percentages over 20% were discarded as well. In total, 21792 genes across 25296 single cells remained for subsequent analysis. The data was natural log transformed and normalized to a total of 1e4 molecules per cell for scaling the sequencing depth by using Seurat. Batch effects were mitigated by using the ScaleData function of Seurat (v2.2.0).

### Identification of cell types and subtypes by dimensional reduction

Seurat (v2.2.0) was used to perform linear dimensional reduction. 931 highly variable genes with an average expression between 0.0125 and 4 and dispersion greater than 1 were selected as input for principal component analysis (PCA). Then we determined significant PCs using the JackStrawPlot function. The top ten PCs were used for t-Distributed Stochastic Neighbor Embedding (tSNE) to cluster cells by the FindClusters function with Resolution 2.0. Clusters were identified by the expression of known cell-type markers and GO analysis.

### Identification of differentially expressed genes among clusters

The differentially expression genes (DEGs) of each cluster were identified by the FindAllMarkers function (thresh.use =0.25, test.use = “bimod”) using the Seurat R package ^89^. The Bimod test returns likelihood-ratio tests for single cell gene expression, and genes with average expression difference > 0.25 natural log with p<0.05 were selected as marker genes. Enriched gene ontology terms of marker genes were identified using DAVID6.7 ^90,91^ (https://david.ncifcrf.gov/home.jsp).

### Integration of the datasets between retina progenitor cells and different types of cells

We setup two Seurat objects belong to retina progenitor cells and different types of retina neurons and müller glia, we remove Ki67 positive cells for further analysis to avoid the cell cycle effects. Then we performed the FindVariableGenes function to detect variable genes in each dataset independently. We take the union of the top 1000 variable genes from both datasets for canonical correlation analysis, and aligned the resulting canonical correlation vectors (CCV) across datasets with the AlignSubspace function (https://satijalab.org/seurat/immune_alignment.html)^66^. tSNE was used for downstream analysis with the top 20 CCV.

### Single cell trajectory analysis

The Monocle2 R package (version 2.6.1) was used to construct single cell pseudo-time trajectories to discover developmental transitions ^54-56^. We used highly variable genes identified by Seurat to sort cells in pseudo-time order. The actual gestational time of each cell tells us which state of cells are at beginning of pseudo-time in the first round of “orderCells”. We then called “orderCells” again, passing this state as the root_state argument. “DDRTree” was applied to reduce dimensional space, and the minimum spanning tree on cells was plotted by the visualization functions “plot_cell_trajectory” or “plot_complex_cell_trajectory”. BEAM tests were performed on the first branch points of the cell lineage using all default parameters. Plot_genes_branched_pseudotime function was performed to plot a couple of genes for each lineage.

### Cell-cycle analysis

In the cell-cycle analysis, we used a cell-cycle related gene set with 43 genes for S phase and 54 genes for G2/M phase of the cell cycle ^92,93^. We assigned each cell a score based on its expression of G2/M and S phase genes by the CellCycleScoring function using the Seurat R package. These gene sets should be anticorrelated in their expression levels, and cells expressing neither G2/M or S phase genes are likely not cycling and in G1 phase.

### Statistics

All data were represented as the mean ± SEM. Comparisons between two groups were made using *t*-tests. The quantification graphs were analyzed using GraphPad Prism (GraphPad Software).

---

**Supplementary Figure 1.**
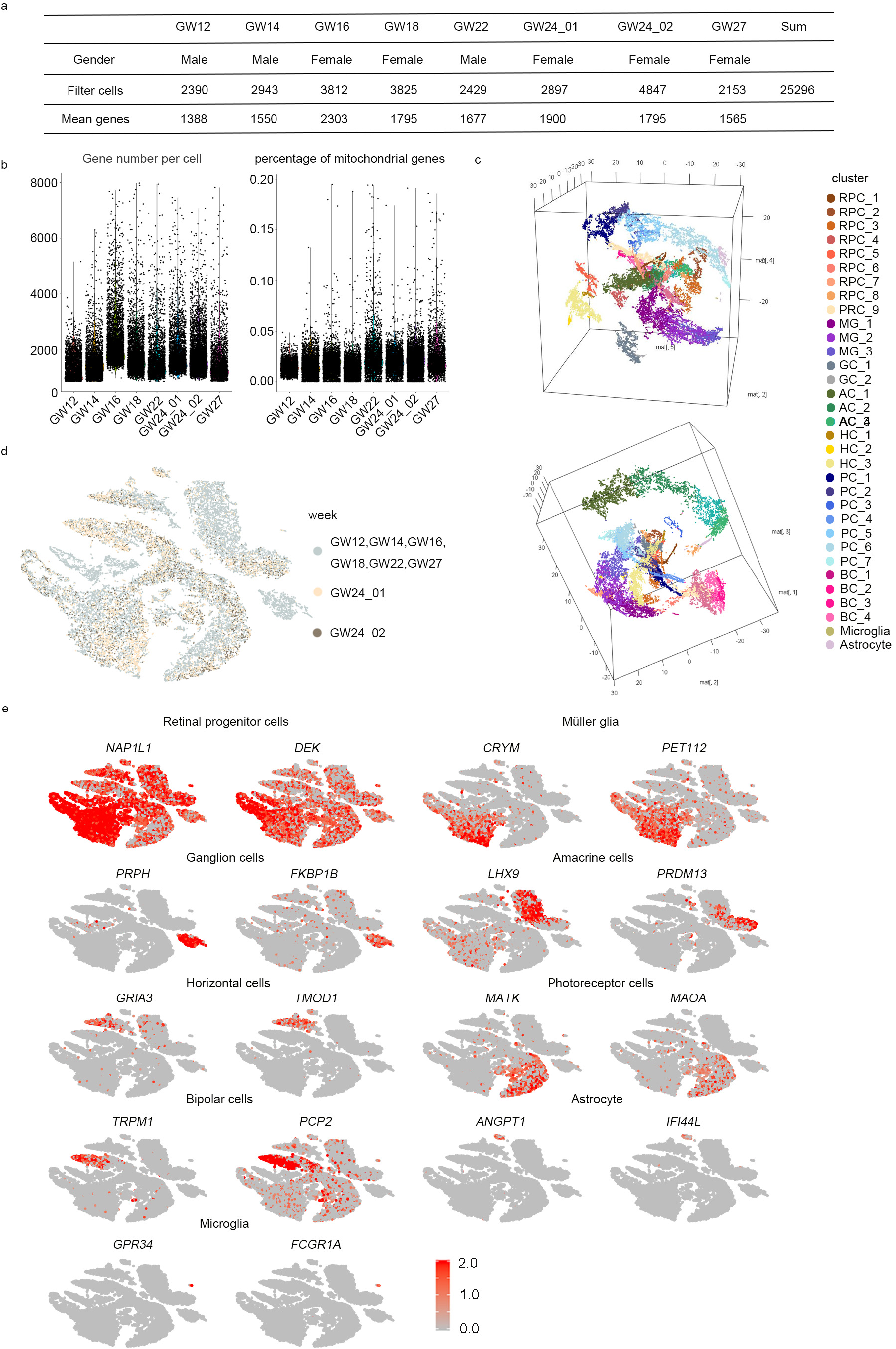
Single-cell RNA-seq Information and Molecular Diversity of the Developing Human Retina. a. Human embryo retinal sampling information.
b. Quality control for eight samples, each dot represents a single cell.
c. 3D t-SNE plots for human embryonic retina. Dimensions 2,4,5 (left top); dimensions 1,2,3 (left bottom). Each dot represents a single cell, and clusters are color-coded
d. t-SNE plots for human embryonic retina. The two repetitions of GW24 are labeled in different colors, and no obvious differences in distribution are observed among the different batches from the same embryo stage. The color represents the gestational week of the sample from which the cell was collected.
e. t-SNE plots of identified novel markers for each cell type.

**Supplementary Figure 2.**
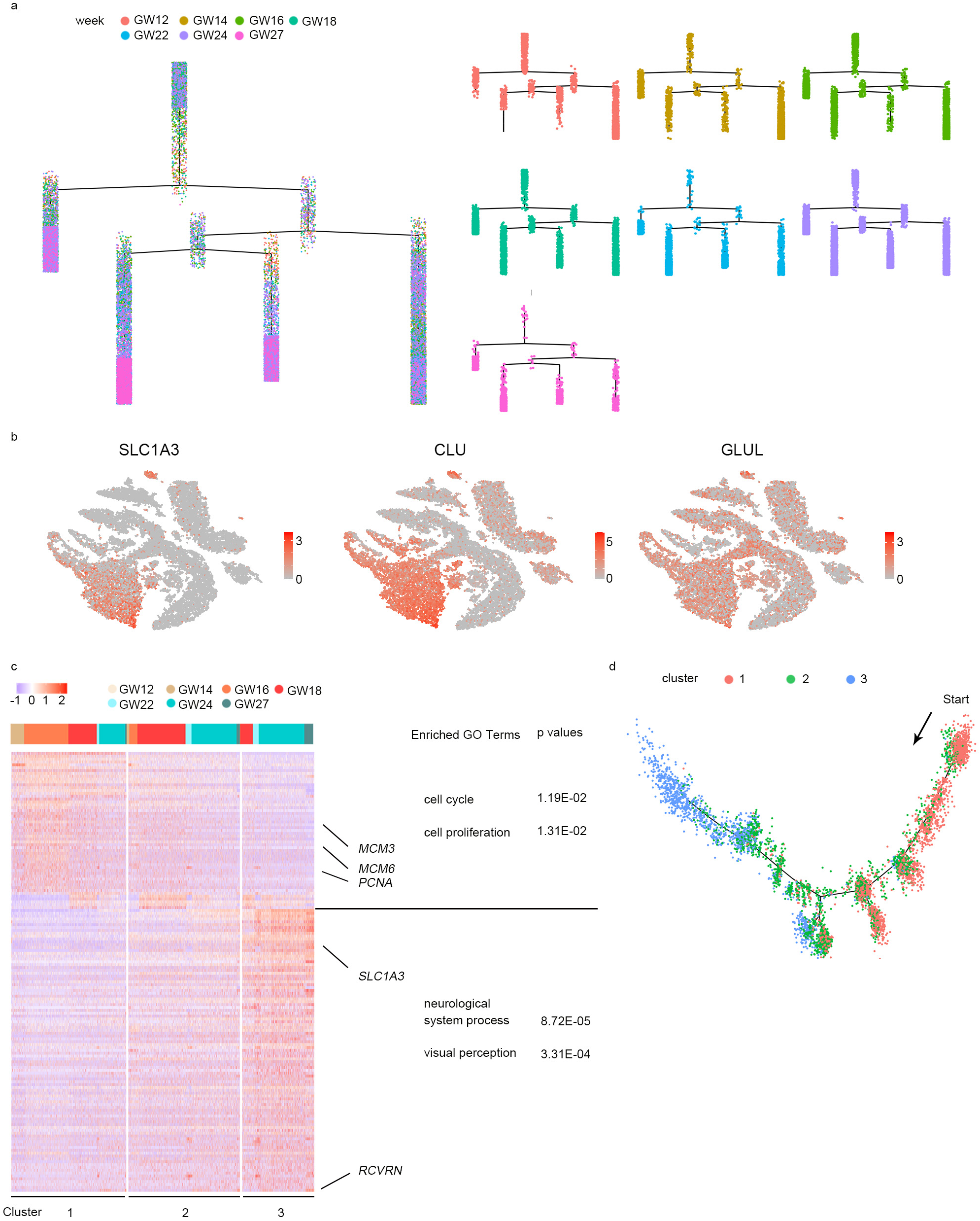
Monocle Analysis of the Developing Human Retina and Subclusters of Müller Glia. a. Single-cell trajectories by Monocle analysis showing the development of the human embryo retina. Each dot is a single cell, and its color represents the gestational week at which it was collected.
b. Expression patterns of known markers for Müller glia on t-SNE plots. Grey to red indicates a gradient from low to high gene expression.
c. Heat map showing the gene expression of Müller glia subclusters. The enriched gene ontology terms are highlighted on the panel to the right. The bar chart on the top shows the gestational week. Blue to red indicates a gradient from low to high gene expression.
d. Cell lineage relationships of Müller glia from 3 subclusters.

**Supplementary Figure 3.**
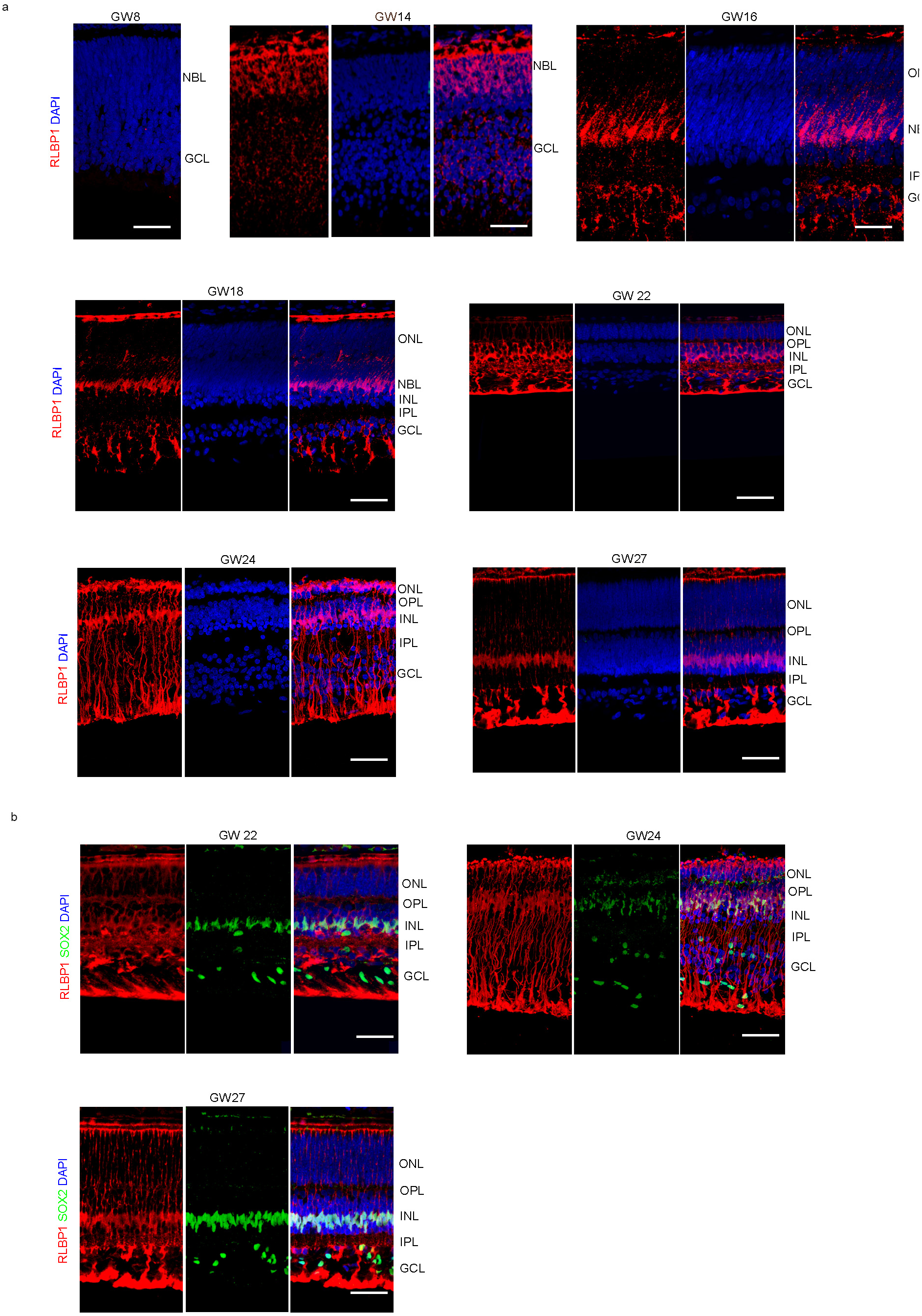
Immunostaining of Müller Glia in the Developing Human Retina. a. Immunostaining with Müller glia marker RLBP1(red) of GW8, GW14, GW18, GW22, GW24 and GW27 samples, showing the position and morphology of Müller glia cells in the human embryonic retina, scale bar 50 μm. Higher-magnification views of regions boxed for Müller glia at GW14 are on the bottom, scale bar 10 μm.
b. Immunostaining with RLBP1(red, Müller glia) and SOX2(green, retinal progenitor cells) in GW22, GW24, and GW27 human retina showing that Müller glia also express retinal progenitor cell markers. Scale bar, 50 μm.

**Supplementary Figure 4.**
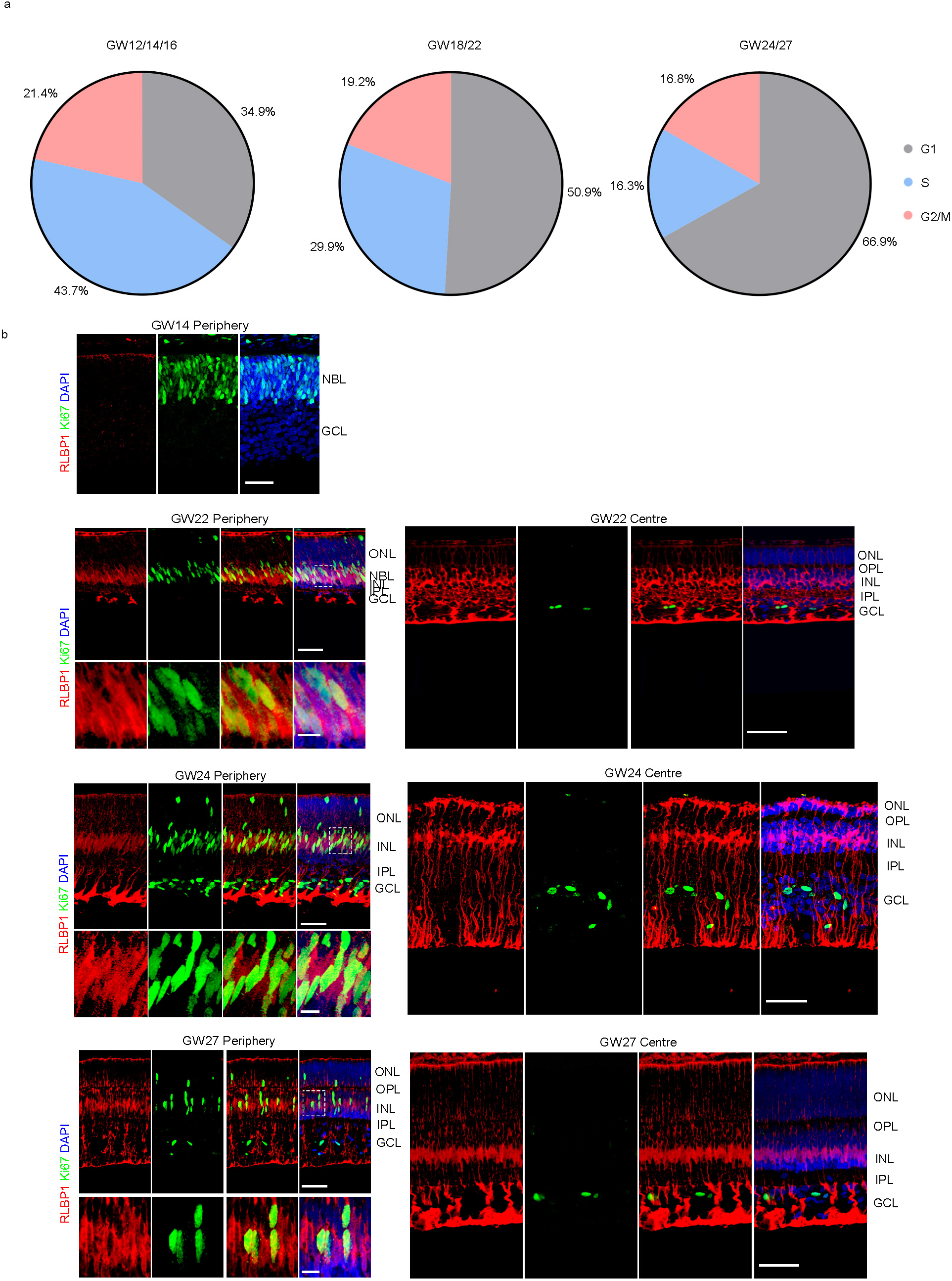
Cell Cycle Properties of Müller Glia in the Developing Human Retina. a. Distribution of G1, S, and G2/M stages of the cell cycle for Müller glia at different gestational weeks.
b. Immunolabelling with RLBP1 (red, Müller glia), Ki67 (green, mitotic cells) at central and peripheral locations of the retina. RLPBP1+ Ki67+ double positive cells were greater in the periphery of the retina compared to the centre. Scale bar, 50 μm(left top, right), 10μm(left bottom).

**Supplementary Figure 5.**
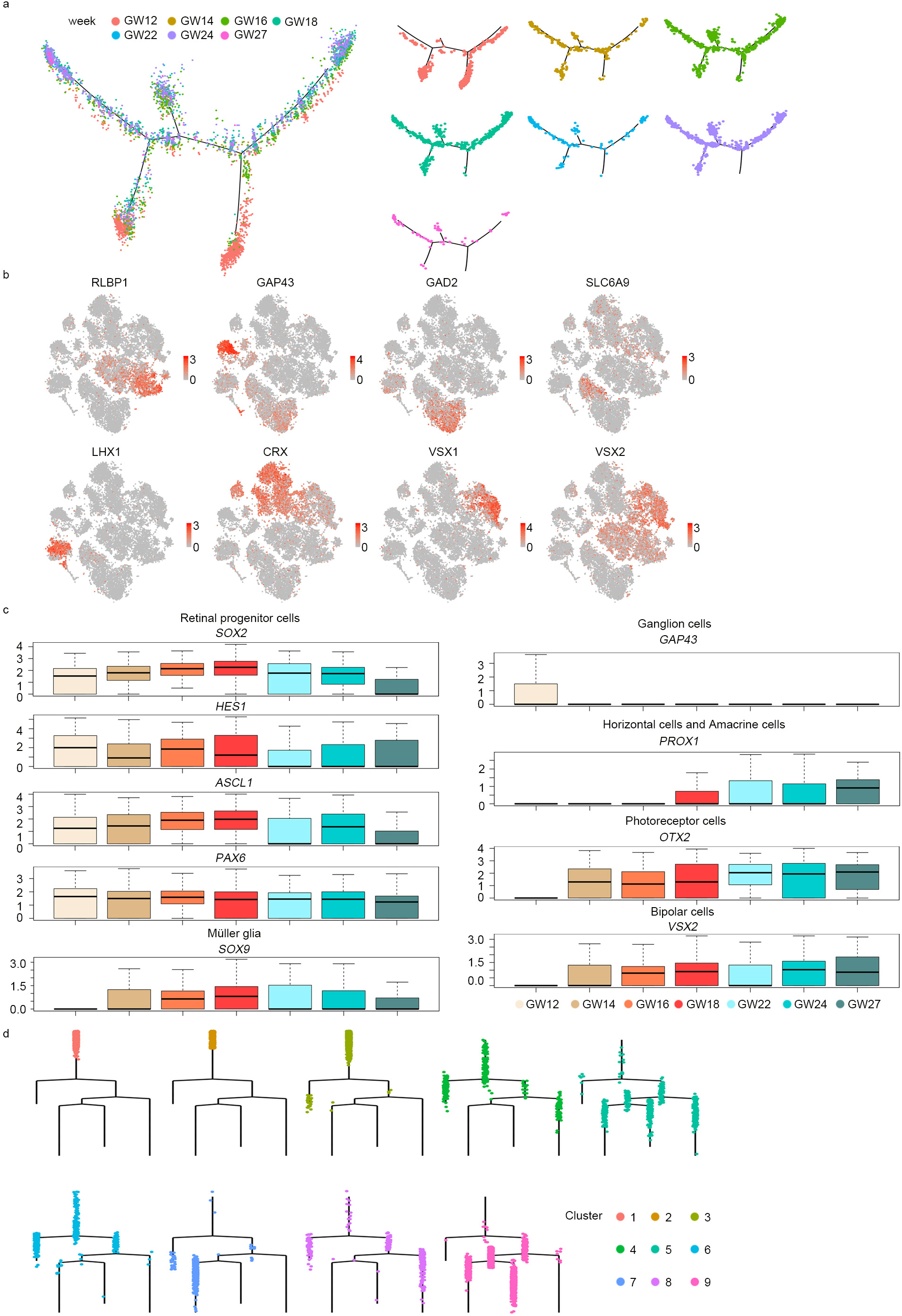
Gene Expression Features of Retinal Progenitor Cells in Human Fetal Retina. a. Single-cell trajectories by Monocle analysis showing the development of retinal progenitor cells. Each dot is a single cell, and its color represents different gestational weeks.
b. Expression patterns of known markers for different cell types on t-SNE plots after Seurat alignment procedure (grey, no expression; red, increased relative expression).
c. Gene expression of known markers for retinal progenitor cells, Müller glia, ganglion cells, amacrine cells, horizontal cells, photoreceptor cells, and bipolar cells found in retinal progenitor cells. The color for box plot shows different gestational weeks.
d. Single-cell trajectories of retinal progenitor cells from different subclusters. Each dot is a single cell, and its color represents the different subclusters. The trajectories layout is based on Monocle analysis in Fig. 2a.

**Supplementary Figure 6.**
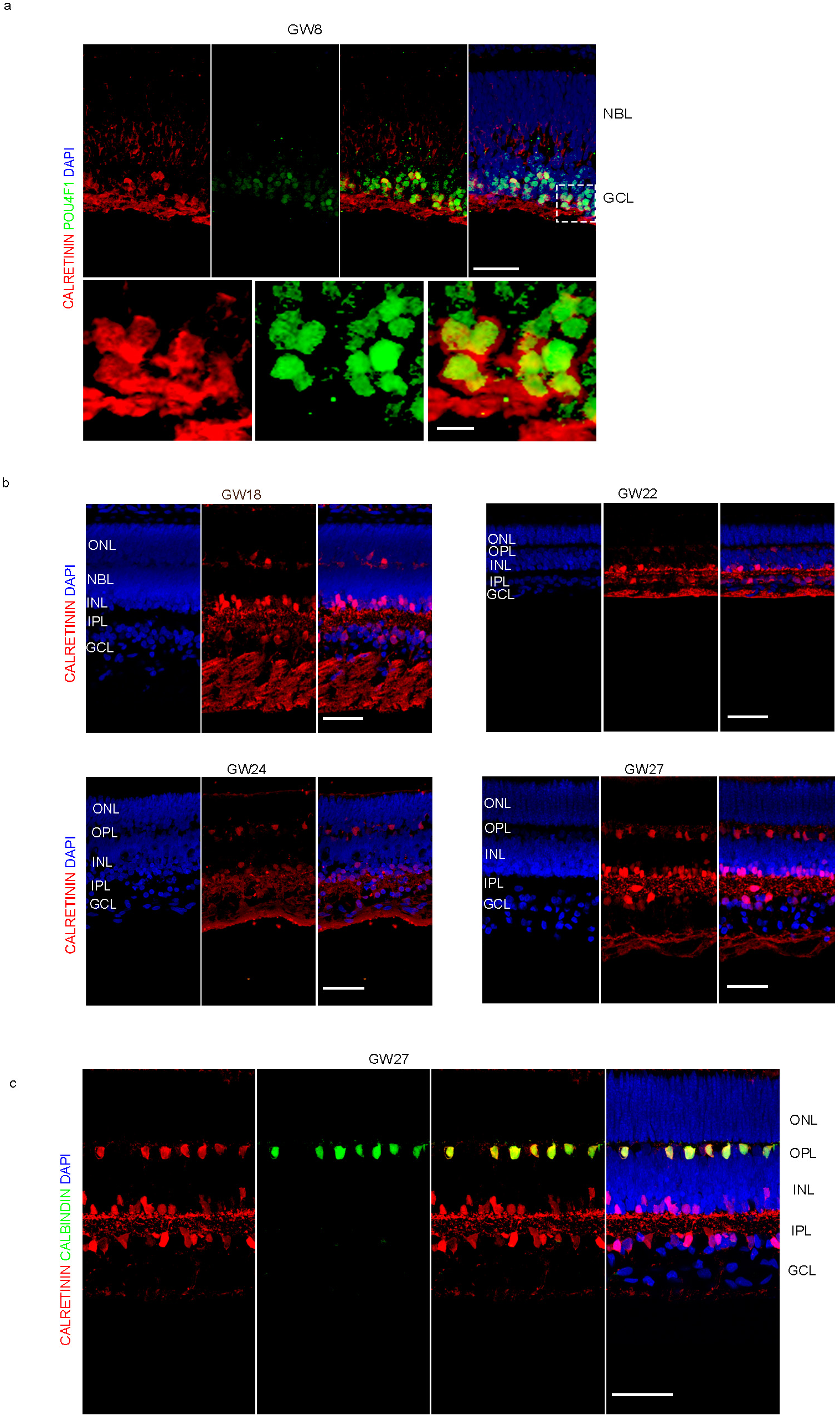
Ganglion, Amacrine and Horizontal Cell Development in the Developing Human Retina. a. Immunostaining with CALRETININ (red, amacrine cells, horizontal cells, ganglion cells) and special ganglion cell marker POU4F1 (green) in the human retina at GW8. All CALRETININ+ cells also express POU4F1, identifying them as ganglion cells. Scale bar, 50 μm(top), 10 μm(bottom).
b. Immunostaining with CALRETININ (red, amacrine cells, horizontal cells, ganglion cells) at GW18, GW22, GW24, and GW27. Scale bar, 50 μm.
c. Immunostaining with CALBINDIN (red, horizontal cell) and CALRETININ (red, amacrine cells, horizontal cells, ganglion cells) in GW27 retina. Scale bar, 50 μm.

**Supplementary Figure 7.**
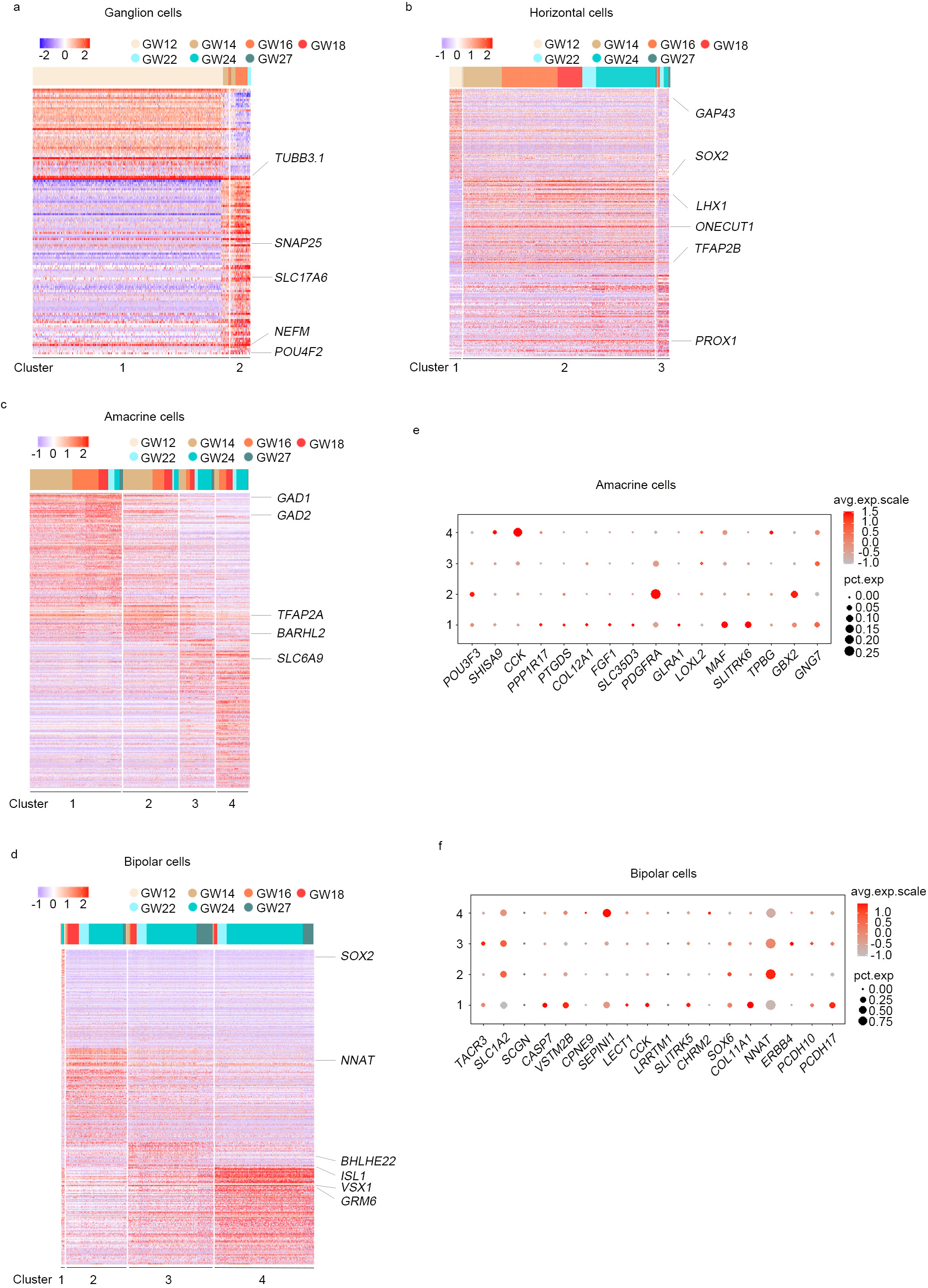
Subclusters of Ganglion, Amacrine, Horizontal and Bipolar Cells. (a-d) Heat map of differentially expressed genes in different subtypes of (a) ganglion cell. (b) horizontal cells, (c) amacrine cells, (d) bipolar cells. Known markers within each subcluster are shown to the right. The graph on the top shows the distribution of each subcluster across gestational week. Blue to red indicates a gradient from low to high gene expression. (e-f) Dot plot for known markers of (e) amacrine cell and (f) bipolar cell subclusters. The size of the dot represents the percentage of cells in the cluster. Grey to red indicates a gradient from low to high gene expression.

**Supplementary Figure 8.**
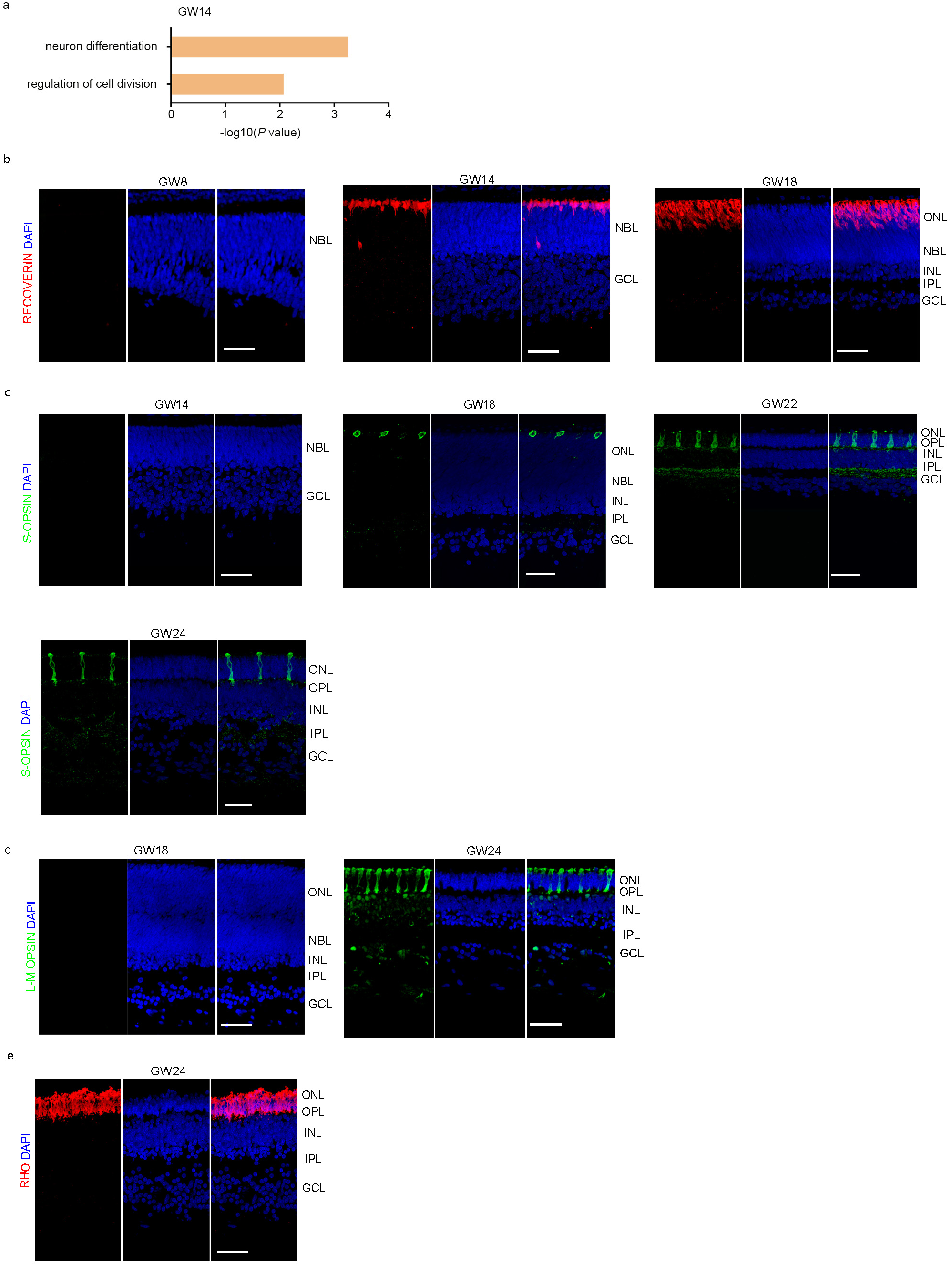
Characterization of Photoreceptor Cell Development in Human Fetal Retina. a. Enriched gene ontology terms show cell properties of photoreceptor cells at GW14.
b. Immunostaining with RECOVRIN (red, early photoreceptor) at GW8, GW14, and GW18. Scale bar, 50 μm
c. Immunostaining with S-OPSIN (green, S-Cone) at GW14, GW18, GW22, GW24, and GW27. Scale bar, 50 μm.
d. Immunostaining with L/M-OPSIN (green, L/M-Cone) at GW18, and GW24. Scale bar, 50 μm.
e. Immunostaining with RHO (red, Rod photoreceptor) at GW24 and GW27. Scale bar, 50 μm.

**Supplementary Figure 9.**
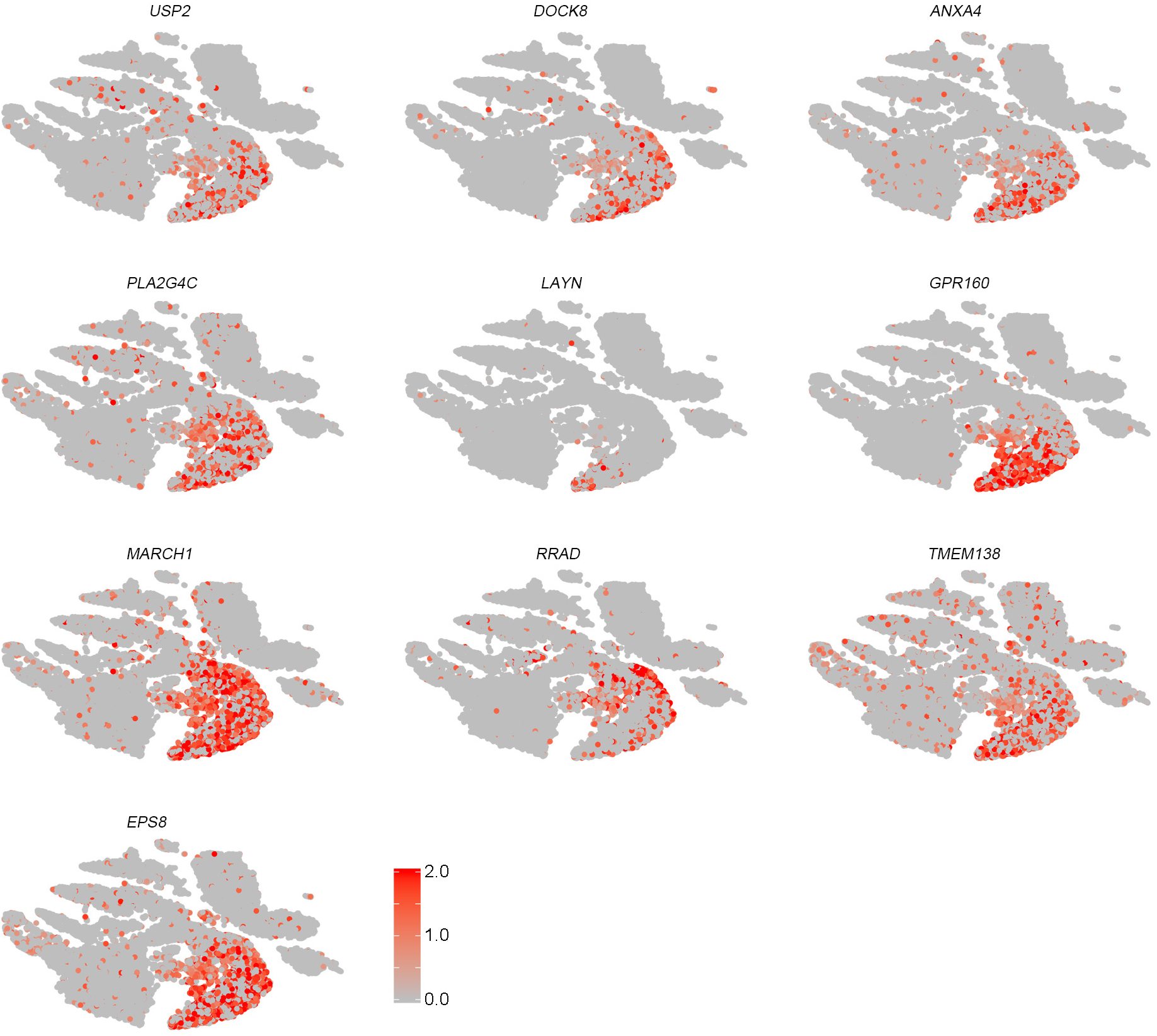
t-SNE plots of identified novel markers for photoreceptor cells (grey, no expression; red, relative expression).

